# Deciphering the metabolic capabilities of Bifidobacteria using genome-scale metabolic models

**DOI:** 10.1101/659888

**Authors:** N. T. Devika, Karthik Raman

## Abstract

Bifidobacteria, the initial colonisers of breastfed infant guts, are considered as the key commensals that promote a healthy gastrointestinal tract. However, little is known about the key metabolic differences between different strains of these bifidobacteria, and consequently, their suitability for their varied commercial applications. In this context, the present study applies a constraint-based modelling approach to differentiate between 36 important bifidobacterial strains, enhancing their genome-scale metabolic models obtained from the AGORA (Assembly of Gut Organisms through Reconstruction and Analysis) resource. By studying various growth and metabolic capabilities in these enhanced genome-scale models across 30 different nutrient environments, we classified the bifidobacteria into three specific groups. We also studied the ability of the different strains to produce short chain fatty acids, finding that acetate production is niche- and strain-specific, unlike lactate. Further, we captured the role of critical enzymes from the bifid shunt pathway, which was found to be essential for a subset of bifidobacterial strains. Our findings underline the significance of analysing metabolic capabilities as a powerful approach to explore distinct properties of the gut microbiome. Overall, our study presents several insights into the nutritional lifestyles of bifidobacteria and could potentially be leveraged to design species/strain-specific probiotics or prebiotics.

## INTRODUCTION

The human gastrointestinal tract harbours a diverse and complex microbial ecosystem, which regulates multiple physiological processes that play a fundamental role in the well-being of their host. This microbiome is associated with a plethora of functions, which include fermentation and absorption of complex carbohydrates, maturation and normal development of immune functions, and prevents adhesion of pathogens to the intestinal surface^1^. Several factors, starting from the mode of delivery, to breastfeeding, gender, age, geography, disease, drug usage and long-term dietary intake, influence the structure and activity of the trillions of microorganisms inhabiting the gastrointestinal tract^2^. Many diseases, notably obesity, coronary heart disease, diabetes and inflammatory bowel disease, have all been associated with dysbiosis in gut microbiota composition^3^. Thus, the gut microbiome is considered a complex polygenic trait, shaped by both environmental and genetic factors^4^.

Bifidobacteria, most frequently isolated from the faeces of breast-fed infants, are involved in the maintenance of intestinal microbial balance and health. Bifidobacteria exert their biological activities through the production of vitamins and antimicrobial substances; further, they regulate the immune system and have anti-obesity and anti-inflammatory activities^5,6^. Bifidobacteria are gram-positive, anaerobic, and saccharolytic, and have been reported to inhabit the intestinal tract of mammals and insects, the human oral cavity and sewage^7^. This genus encompasses a broad range of enzyme-coding genes associated with the uptake and catabolism of complex and non-digestible carbohydrates, ranging from human milk oligosaccharides to plant fibre^8^. Bifidobacteria degrade the hexose sugars glucose and fructose through a unique pathway named “bifid shunt”, which is centred on the key enzyme fructose-6-phosphate phosphoketolase^9^. The metabolites from this ATP-generating pathway mainly produce short-chain fatty acids (SCFAs) that antagonise pathogenic bacteria and form a barrier against infection^10^. For instance, acetate produced by bifidobacteria improves intestinal defence mediated by epithelial cells, and thereby protects the host against lethal infections^11^.

Several species of bifidobacteria, namely *B. animalis, B. breve* and, *B. longum* are used to treat various gastrointestinal disorders and inflammatory bowel disease^12^. Also, a few strains of bifidobacteria such as *B. animalis* BF052, and *B. animalis subsp. lactis* BB-12 form the major functional ingredients in commercialised probiotic food products^13^. Notably, the probiotic characteristics of bifidobacteria are strongly strain-dependent^14^, with applications in the food, dairy, and pharmaceutical industries. Therefore, an understanding of the metabolism of this genus in its entirety and the adaptation of distinct strains to a variety of nutrient environments is definitely necessary.

In the recent decade, considerable effort has been invested into understanding the gut microbiome using metabolic modelling, particularly constraint-based reconstruction and analyses, which generate testable hypotheses to elucidate the metabolism of individual species, and also interspecies metabolic interactions^15–17^. Recently, Thiele and co-workers^18^ generated AGORA (Assembly of Gut Organisms through Reconstruction and Analysis), an excellent resource of semi-curated genome-scale models specifically for human gut microbes, enabling system-level studies of the gut. These genome-scale metabolic models of gut microbes have been used to predict growth phenotypes under different conditions and also provide a link between dietary intake and absorption in humans^19,20^. Moreover, research on strain-specific metabolic reconstruction has been explored for strains of *E. coli*^21^, *Staphylococcus aureus*^22^ and *Salmonella*^23^. These studies exemplify the use of metabolic networks to probe strain/species-specific diversity, and provide insights into the utility of different strains towards their diverse applications.

The identification and classification of *Bifidobacterium* species have been demonstrated using DNA-DNA hybridization^24^, and by creating a phylogeny based on whole and/or conserved genomic sequences^25^. In the present study, we set out to capture the diversity between strains of bifidobacteria by generating condition–specific metabolic models with the genome-scale metabolic models obtained from AGORA and subsequently investigating various phenotypic and metabolic characteristics. We report pronounced differences across strains with respect to the nutrient utilisation, metabolic capabilities, variability in the bifid shunt pathway, and essential reactions under diverse niches. Ultimately, our modelling approach enabled us to classify the bifidobacteria into three groups, i) *B. bifidum*, ii) *B. animalis*, and iii) *B. longum* based on multiple phenotypic and metabolic properties. In summary, this study employs a multi-pronged modelling approach to systematically characterise bifidobacteria based on their various phenotypic and metabolic features under different nutritional environments.

## RESULTS

In this section, we illustrate how our constraint-based modelling approach enables a careful classification of 36 strains of *Bifidobacterium* and contributes to a detailed understanding of their metabolism and metabolic capabilities. Our key results are four-fold. First, we show that many of our (re-curated) individual models better capture experimental observations in metabolising multiple carbon sources. Second, we analyse strain-specific SCFA production under 30 different nutrient environments and illustrate the key differences between the metabolic capabilities of different bifidobacteria. Third, we distinguished the strains further, based on the reaction knock-outs performed in the bifid shunt pathway, across strains and in different environments. Together, these approaches provide a firm basis to understand the metabolic diversity of bifidobacteria and their roles in maintaining gut health. Next, we computed single lethals in a rich environment and identified the essential reactions across strains. Finally, our studies unravel three distinct clusters of these 36 strains, based on multiple phenotypic and metabolic properties.

### Bifidobacteria cluster into two groups based on carbohydrate utilisation

Based on the phenotypic prediction with respect to carbohydrate utilisation, the 36 strains could be differentiated into two groups, as shown in Fig 1. All strains included in this study exhibited growth on glucose, fructose and maltose. The strains BGN4 and NCIMB 4117 demonstrated limited growth compared with other species of *Bifidobacterium*. Further, among the strains studied, *B. adolescentis* ATCC 15703 could utilise most of the carbon sources used in this study, making the strain nutritionally versatile, followed by *B. dentium* ATCC 27678 and *B. kashiwanohense* DSM 21854 belonging to *B. adolescentis* group^25^. The probiotic strain *B. bifidum* BGN4 was the only strain that could not utilise the prebiotic carbohydrate inulin. In addition, we could observe that *B. longum infantis* ATCC 15697 differed from all of the other strains from this same species in fermenting mannitol and trehalose and not fermenting arabinose and arabinotriose. The widely used commercial species, namely *B. animalis,* clustered together, with *B. animalis* BB 12 showing a distinct ability to ferment mannose. Overall, we observed distinct substrate utilisation profiles for our diverse collection of bifidobacterial strains.

**Fig 1:**
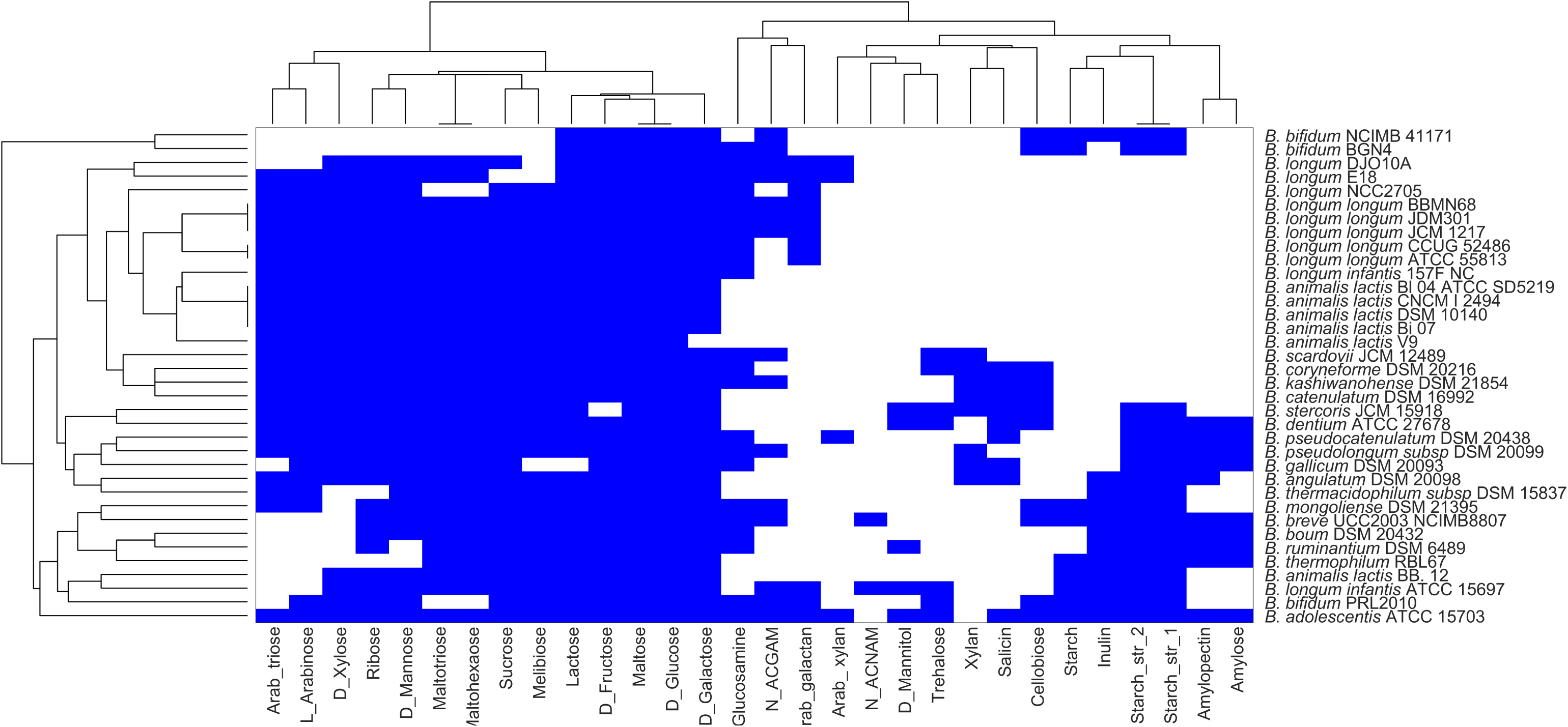
Clustering of bifidobacterial species grown on multiple nutrient environments. Rows represent individual strains and columns represent different nutrient environments. Strains are clustered based on the ability to sustain growth in each of the different nutrient environments. The presence and absence of growth are indicated by blue and white colours, respectively.

### Lactate and acetate production vary across strains in different environments

To determine the metabolic capabilities of bifidobacteria, we performed Flux Variability Analysis (FVA; see Methods), and the production of metabolites, namely, acetate, lactate, ethanol, succinate, and formate, were assessed. FVA essentially indicates the possible range of variation for the flux of every reaction, under specified conditions (e.g. maximum growth).

Based on the clustering of the synthesis profiles of SCFAs in diverse nutrient environments, three major divisions can be distinguished among the bifidobacterial strains, as shown in Fig. 2. Among the SCFAs, the present study majorly focussed on acetate and lactate, which make a significant contribution to the prevention and treatment of metabolic syndromes^26^. Among the studied strains, *B. kashiwanohense* DSM21854, *B. thermophilum* RBL67, *B. gallicum* DSM 20093, *B. longum infantis* 157F NC and *B. pseudolongum subsp. pseudolongum* DSM 20099, were found to be producers of acetate as well as lactate in their survivable environments.

**Fig 2:**
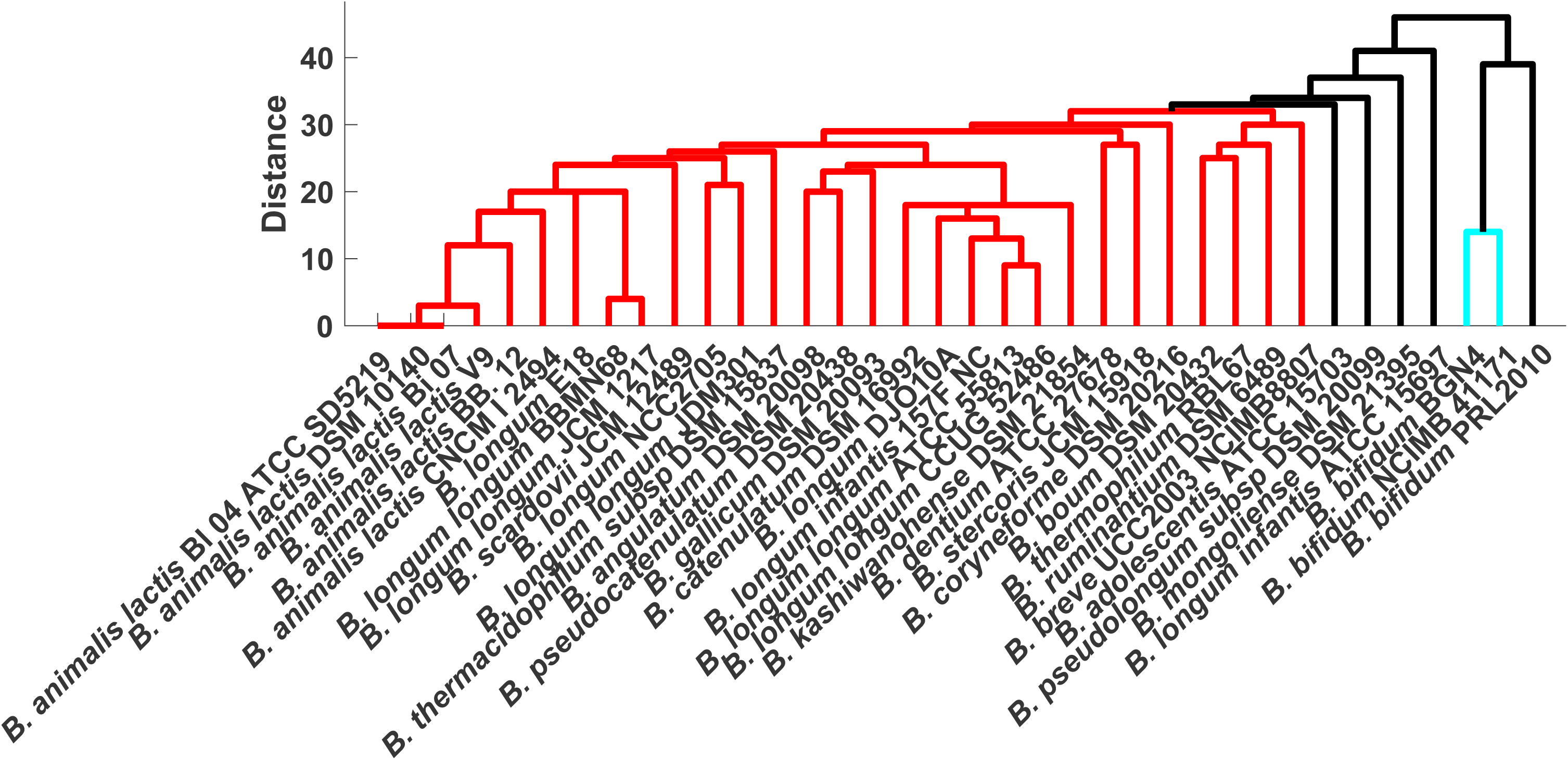
Dendrogram representation of bifidobacteria based on SCFA production. SCFA production of strains under diverse nutrient environment was represented as binary vectors and used to calculate the Hamming distance between strains.

However, lesser acetate production was associated with CNCM I 2494 strain of *B. animalis*. A strain-specific difference was exhibited among *B. bifidum*, with the strain *B. bifidum* PRL 2010 showing a comparatively higher potential for acetate production. The strain *B. bifidum* PRL 2010 also clustered with the nutritionally versatile strain *B. scardovii* JCM 12489. In contrast, *B. adolescentis* ATCC 15703, the strain which utilised most of the carbon sources could not produce acetate in all its survivable environments.

In contrast to acetate production, nearly all the bifidobacterial strains could produce lactate across their survivable nutrient environments. The exceptions were two strains of *B. longum,* namely NCC 2705 and JDM 301, which lack lactate production when monosaccharides were used as the sole nutrient source, with a clear shift of flux towards formate production. Possibly, the assigned carbon uptake was found to be limiting for these strains of *B. longum*. Unexpectedly, the strains with lesser fermentation ability, i.e. the strains that show growth on fewer carbon sources (Fig 1), namely, *B. bifidum* BGN4, *B. bifidum* NCIMB 41171 and *B. bifidum* PRL2010, produce acetate and lactate in all (growth supporting) environments. Notably, among the nine probiotic candidates included in this study, only, *B. thermophilum* RBL67 was able to produce the major metabolites, namely acetate and lactate in all (growth supporting) environments. Further, we could observe that polysaccharides, namely starch, inulin, maltohexaose, arabinotriose and arabinoxylan, displace the flux of *Bifidobacterium* strains towards lactate production. But with monosaccharides, a balance of acetate and lactate production is clearly evident (Supplementary Fig 1).

We further explored the metabolic capabilities of *Bifidobacterium* strains in a carbon rich environment, by allowing uptake for all mono-, di-, tri-, and polysaccharides separately. As observed in the individual nutrient source utilisation, *B. animalis* CNCM I 2494 strain could produce acetate when allowed an uptake for all monosaccharides considered in the study and not with di-, tri-, or polysaccharides. In the case of *B. longum* strains namely, NCC 2705, and JDM 301, allowing uptake of all monosaccharides together resulted in lactate production. This concurs with the results mentioned above where the uptake of a single monosaccharide was found to limit lactate production. Taken together, our analyses show that the production of acetate turns out to be influenced by the nutrient environment and varies across strains. On the other hand, lactate production was independent of the nutrient environment.

### How dispensable is the bifid shunt pathway in *Bifidobacterium* across environments?

Fundamentally, central carbon metabolism contributes to the synthesis of precursor molecules necessary for the growth and survival of an organism. Our analysis focuses on the bifid shunt pathway, a unique and effective central fermentative pathway in the genus *Bifidobacterium*. This glycolytic bypass pathway plays a central role in secreting the SCFAs lactate and acetate, which are important for hosts not only in preventing the growth of harmful bacteria by lowering intestinal pH, but also serve as an energy source for intestinal epithelial cells. We simulate all possible reaction knock-outs in the bifid shunt pathway to understand how dispensable the reactions are, across bifidobacteria.

To this end, we compared 26 different reactions associated with the bifid shunt pathway across strains (Supplementary Table 1), and the reactions were categorised based on their ability to carry flux under different nutrient environments. Intriguingly, only seven reactions turned out to be carrying flux across all environments, and 19 reactions were found to be environment-specific. We designate such reactions as *conserved* and *non-conserved*, respectively. We next performed a reaction knockout on the seven conserved reactions one at a time in each of the environments. We identified three reactions namely, phosphoglucomutase (PGMT), phosphoglycerate mutase (PGM), and phosphoglycerate kinase (PGK) with a major impact, and ribulose 5-phosphate 3-epimerase (RPE) with a lesser impact on the viability of strains. Deletion of the reactions PGK, PGM and PGMT abolish viability in all strains of *B. animalis* and other strains, namely *B. coryneforme* DSM 20216, *B. longum longum* JDM 301, and *B. gallium* DSM 20093 across all nutrient environments (Fig 3). Among the nine probiotic strains analysed, only *B. thermophilum* RBL 67 was viable upon all three reaction knockouts performed across nutrient environments.

**Fig 3:**
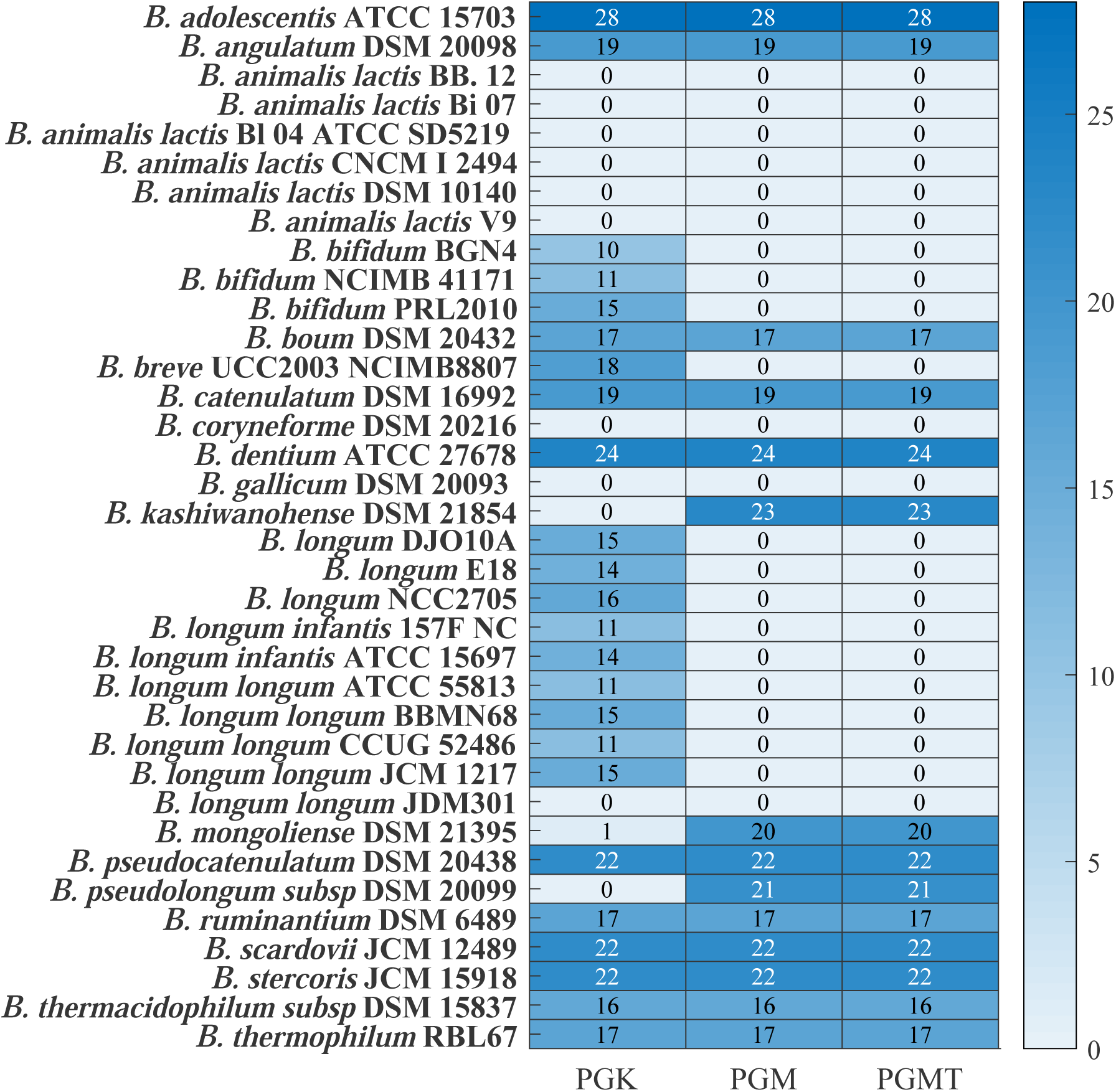
Overview of growth outcomes upon deletion of PGK, PGM, and PGMT in all 30 different nutrient environments. The rows represent 36 strains of bifidobacteria. The colour bar represents the number of environments in which the organisms can survive upon deletion of PGK, PGM and PGMT individually.

Thus, PGK, PGM, and PGMT are essential for nine strains across environments. Further, in our analysis, we identified the reactions PGM and PGMT to be essential for all strains of *B. longum* and *B. bifidum.* Taken together, we identified a set of species-specific environment-independent essential reactions. Further, deletion of these enzymes resulted in production of acetate in more environments and lactate production in fewer environments in the surviving strains of bifidobacteria. This is expected because deletion of either of these enzymes lowers the NADH availability, which in turn reduces lactate production.

In contrast, we found that many other strains of bifidobacteria bypass these reactions and can sustain growth across environments. Therefore, we investigated the corresponding reaction pairs that maintain viability for these strains. Here, we applied the concept of synthetic lethality to identify the reaction pair that rescued growth in many strains. Synthetic lethality analysis was performed using the Fast-SL algorithm (see Methods) derived single and double reaction lethals. Interestingly, it was clearly evident that, in other strains that bypass these reactions, these enzymes were found to be double lethal (Supplementary Table 3), where the organism sustains growth with acetate and lactate production.

### Reaction essentialities in a rich environment reveal diversity between strains

Following the classification of bifidobacterial based on nutrient utilisation and metabolic capabilities, we aimed to obtain a better understanding of the essential reactions across strains. The mammalian intestine, which the bifidobacteria colonise, collectively possesses a wide range of carbohydrates. Hence, we simulated a rich environment by supplementing all the 30 different nutrient sources. We here identify reactions essential for the survival of the genus *Bifidobacterium* and analyse to what extent this essentiality is conserved across species/strain. Among the set of organisms used in the study, *B. ruminantium* DSM 6489 and *B. gallicum* DSM 20093 carry the maximum and the minimum number of essential reactions, respectively (Supplementary Table 3).

We find 459 reactions (excluding exchange reactions) to be essential across strains of bifidobacteria. Out of these 459 reactions, 169 were found to be the *core set* of essential reactions, that is, they were present in all the 36 strains. Next, we investigated in which biological pathways these essential reactions function. As expected, most of the essential reactions significantly fall in pathways that synthesise cell wall components, amino acids metabolism (phenylalanine, valine, leucine and isoleucine) and purine synthesis. The grouping of the strains by the effect of single lethal reactions revealed two major classifications of *Bifidobacterium.* The ten strains of bifidobacteria from Group 1 (Fig 4) carry fewer single lethal reactions compared to the total essential reactions of other strains.

**Fig 4:**
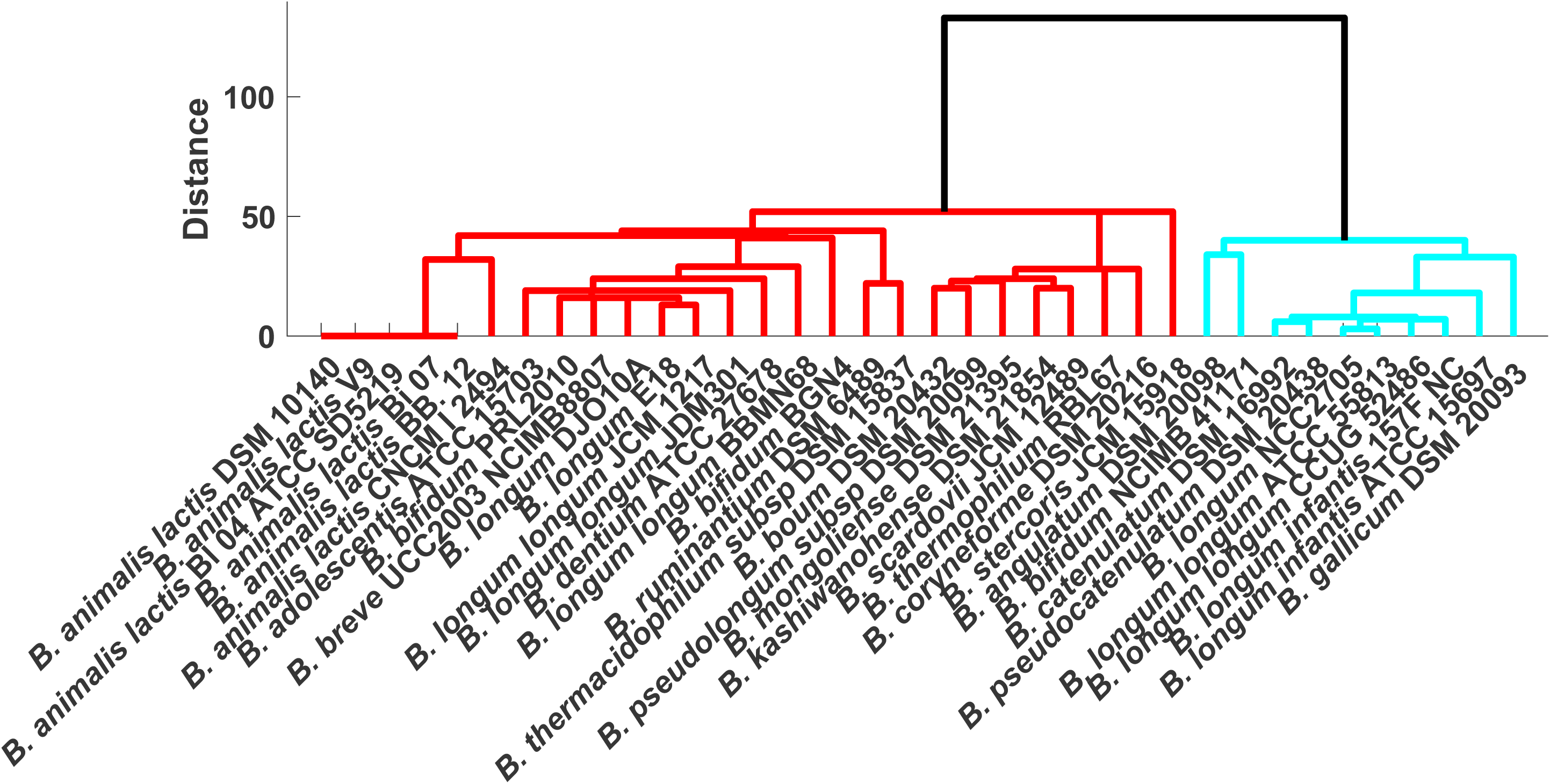
Dendrogram representation of Bifidobacterial species based on the presence of single lethals. Single lethals generated with Fast-SL in the rich environment were grouped based on the Hamming distance between strains.

## DISCUSSION

Bifidobacteria are the quintessential gut inhabitants, which modulate the metabolic and immune activities of their host^27^. Their ability to utilise a wide variety of complex substrates reflects their metabolic adaptations^28^. Studies reveal that bifidobacteria exert strong effects in shaping the bacterial gut community by secreting the major fermentation end-products, such as acetic acid and lactic acid^29^. Moreover, *Bifidobacterium* (including *B. animalis subsp. lactis, B. longum, B. adolescentis, B. breve* and *B. bifidum*) have also been reported to effectively relieve metabolic disorders^30^. As distinct strains showcase different capabilities, it is imperative to understand the inter- and intra-strain diversity between species.

Even though 16S-rRNA sequencing and DNA–DNA hybridisation have been demonstrated to be useful techniques for discriminating bacterial species, these techniques could not provide sufficient resolution to discriminate between closely related bacterial strains^24^. It is well-known that even closely related strains vary in their metabolic repertoire^31^. Therefore, in the present study, emphasis has been laid on genome-scale metabolic networks, as they elucidate the differences with respect to the metabolic capabilities of the organisms. The genome-scale metabolic networks used in this study^18^ represent the entire metabolic repertoire of *Bifidobacterium*. These metabolic models were further curated, to enable the simulation of condition-specific models with a defined set of universal media components.

We began our analysis by computing growth phenotypes on 30 different nutrient environments to determine the distinction between strains. The species commonly identified in the human adult intestine, such as *B. adolescentis,* are enriched in genes that are predicted to be involved in the metabolism of complex carbohydrates^32^. This highlights the metabolic gains made by these species upon adapting to human adult colon^33^. The study also supports the greater genetic diversity exhibited by *B. adolescentis* group that has evolved towards an ecological niche where plant derived saccharides are abundant, such as large intestine of adult human beings^32^. The limited fermentation ability of *B. bifidum* strains correlates with a relatively lower number of carbohydrate transport system compared to other enteric intestinal bifidobacteria^34^. Remarkably, throughout our analysis, all investigated *B. animalis* strains were placed on the same branch in the dendrogram, indicating homogeneity and a high degree of genome conservation in these strains as reported^33^.

Inulin, which occurs naturally in many foods of vegetable origin, is an effective prebiotic as it promotes the proliferation of bifidobacteria^35^. Inulin utilisation by adult and infant strains observed in this study exclude any relationship between strain origin and fermentation ability. Moreover, several bifidobacteria ferment common saccharides, rendering them competitive in the environment. All these analyses reveal the extensive fermentation capabilities of distinct strains of bifidobacteria involved in uptake and utilisation, in particular, polysaccharides such as amylopectin, pullulan, maltotriose, and maltohexaose.

Next, we explored the secretion of SCFAs, particularly lactic and acetic acids, by individual bifidobacteria under different nutrient environments. These SCFAs were selected because they can protect the host against lethal infection^36^. One of the promising probiotic candidates^37^, *B. thermophilum* RBL67 used in this study can produce acetate and lactate in all its survivable niches. This capability of *B. thermophilum* RBL67 could favourably be one of the contributing factors for the antagonistic and protective effects against *Salmonella* and *Listeria* species^38^. The *B. longum* strains used in this study, namely, NCC2705 and JDM301 failed to produce lactate even with monosaccharides say glucose, with an uptake of 10 mmol/gDW/h. This significantly indicates the lesser carbon availability, which limits lactate production.

Interestingly, comparison of other metabolites, namely ethanol, succinate and formate, reveal a shift in metabolism towards formate production in the above-mentioned two strains of *B. longum*. Similar observations were shared by *B. longum subsp. longum* UCD401, which secrete high concentrations of formate with xyloglucans as substrate^39^. Furthermore, bifidobacterial secretion of formate while utilising glucose has also been previously observed^40^. Studies of Wang *et al*^29^ show the importance of *Bifidobacterium* in relieving constipation mainly by improving the concentration of acetic acid in the intestine. In addition to the role of *B. longum* in acetate production as reported by Wang *et al*, our study could identify *B. gallicum* and *B. kashiwanohense* to be capable of producing acetate in all viable environments. In addition, we were able to establish the possible role of polysaccharides, namely starch, inulin, maltohexaose, and amylopectin, which majorly shift the flux towards lactate production across strains. Overall, the contribution of acetate/lactate production by each strain was found to be variable across nutrient environments, revealing the need to understand the strain-specific differences that could probably give a clue for the selection of the most relevant strains for distinct applications.

We further considered the bifid shunt pathway and studied how variable or conserved are the reactions in the pathway across strains under multiple nutrient environments. We found that the enzymes, namely PGK, PGM and PGMT from the bifid shunt pathway exhibited a major impact on the viability of a subset of strains across all nutrient environments. Metabolism is often viewed as an assemblage of several functional modules that coordinate to perform biological activities. However, given the fact that all strains share the enzymes, PGM and PGK were essential to only a subset of species/strains. This highlights the existence of a unique network structure, where strains take up alternative metabolic routes or compensatory reactions to furnish energy and biomass precursors necessary for growth. These observations also provide insights into the diversity of metabolic reactions across strains of the same genus.

To further investigate the metabolic diversity, we computed essential reactions across the strains in a rich environment. The reaction essentialities enabled us to identify two main clusters of *Bifidobacterium,* reflecting species/strain-specific differences in the metabolic reactions. We identified a core set of 169 reactions, essential to all bifidobacteria—given the central importance of bifidobacteria in human health, it will be important to make sure that new antibiotics do not interfere with this core set of essential reactions.

To summarise, we classify the bifidobacteria into three distinct groups:

**Group 1**: The strains of the species, *B. bifidum*, namely NCIMB 41171 and BGN4 are well separated from all 34 strains of *Bifidobacterium* based on their phenotypic predictions, SCFA production, as well as local differences within the bifid shunt pathway.

**Group 2**: The strains of *B. animalis*, used specifically in the food industry, represent a group with highly similar phenotypic and metabolic characteristics. In particular, it forms a cluster with *B. longum,* based on growth and with local differences in the bifid shunt pathway. On the other hand, it forms a cluster with *B. thermacidophilum subsp. thermacidophilum* DSM 15837 based on the SCFA production.

**Group 3:** The ten strains of *B. longum,* being from same species, however are not identified as one group due to diverse nutrient and metabolic capabilities. The strains namely, *B. longum longum* ATCC 55813, *B. longum longum* CCUG 52486, and *B. longum infantis* 157F NC tends to group based on phenotypic and metabolic characteristics.

Further, the other species of *Bifidobacterium* do not follow any interesting pattern, which could serve as a separate cluster. Our analysis did not find any similarities between species of probiotics significance. Further, no similarities exist between the most prominent infant species, namely *B. breve, B. bifidum*, and *B. longum*. A classification of this sort is necessary as it even helps in distinguishing the closely related microbes such as *Lactobacillus* and *Bifidobacterium* species to maintain a stable co-existence with each other and the host. Moreover, the work also highlights the potential of constraint-based approaches namely FBA and FVA to understand the contribution of individual species/strains towards fermentation and production of metabolites that will help in identifying the relevant strains with diverse applications.

Our study does have its limitations. Firstly, we are limited by the number and quality of the genome-scale reconstructions presently available. Consequently, some of the strains in this study are not as well-represented as others; while we have 6 strains of *B. animalis*, we have only one strain *of B. adolescentis* ATCC 15703. Further, BLASTP homology searches performed do not necessarily ensure that the corresponding enzymes are functional, which needs further experimental validation. Even though bifidobacteria thrive in the large intestine, an anaerobic environment rich in complex carbohydrates, it remains a challenge to consider the complex saccharides like human milk oligosaccharides (HMO), fructo-oligosaccharides (FOS), and galacto-oligosaccharides (GOS). The effect of how these complex saccharides influence the metabolic capabilities of *Bifidobacterium* need to be examined in future studies. As no experimental constraints (e.g. internal flux measurements arising out of ^13^C metabolic flux analysis) were used in the model, it should be noted that the variations observed between distinct strains must be taken in a more qualitative context, rather than quantitatively, in terms of the exact values of the fluxes.

Overall, the present study lays special emphasis on the nutrient utilisation, metabolic capabilities, and bifid shunt pathway of *Bifidobacterium*. With the refined genome-scale metabolic models of bifidobacteria, we anticipate that regardless of whether from the same species, we observed inter-and-intra-strain differences at fermentation and metabolic capability. Taken together, the analyses mentioned above present a comprehensive summary of knowledge regarding the metabolism of *Bifidobacterium*. We believe that the deeper understanding of bifidobacterial metabolism thus obtained will better facilitate the use and exploitation of bifidobacteria in various commercial applications.

## METHODS

### Data

We used the recently generated comprehensive resource of 773 semi-curated constraint-based metabolic models of relevant gut microbes, AGORA^18^. AGORA contains 39 strains of *Bifidobacterium,* of which we chose 36 strains for our analysis. We eliminated three other strains (*B. animalis lactis* AD011, *B. bifidum* S17, and *B. breve* HPH0326), as they produced growth even in the absence of the representative carbon source glucose. The genome-scale metabolic models of *Bifidobacterium* were obtained from Virtual Metabolic Human (VMH) Database, a resource of semi-curated models of gut microbes, AGORA. The 36 strains (covers 20 different species) investigated in this study and the general information on the origin of these species is listed in Supplementary Table 1. Among these twenty species, *B. breve, B. bifidum,* and *B. longum* are the most prominent species in the infant gut^41^. Several of the other species used in this study are even commercially important, often used as a probiotic supplement.

### Model expansion to account for experimental growth profiles

Initially, we focused on *B. bifidum* PRL 2010, a strain that is very well described phenotypically and genotypically. We generated a *condition-specific* model that could recapitulate the previously reported experimental observations under various growth conditions. We simulated a defined set of media components reported in the literature to achieve *in silico* growth for validation of the model. We examined the growth of the *B. bifidum* PRL2010 model on seven different carbon sources, namely glucose, xylulose, fructose, lactose, galactose, sorbitol and maltose. We observed that the model did not demonstrate growth on fructose as the sole carbon source; to address these, we added additional transport reactions for fructose and glucose, expanding the model. Finally, our simulations were consistent with reported experiments, confirming the accuracy of the model in utilising different carbon sources^42^.

We also manually re-curated 35 genome-scale models of other bifidobacteria to improve their fit to experimental observations in utilising the different carbon sources (Supplementary Table 2). During the course of our analyses, most of the models failed to show growth on carbon sources where growth has been demonstrated in previous experiments. Except *B. adolescentis* ATCC 15703, none of the AGORA models demonstrated growth in the presence of starch. However, it is well-documented that most strains of bifidobacteria do hydrolyse starch^43^. The AGORA models failed to capture such starch utilisation, as most of the models lack reactions pertaining to starch metabolism. Therefore, we carried out a standalone BLASTP (with an E-value cut-off of 10^-5^) to identify putative enzymes associated with starch degradation across different strains. We observe that 31 strains of bifidobacteria do carry a homologue of the enzymes oligo-1, 6-glucosidase, *α*-1,6-pullulanase and *α*-1,4-amylase associated with starch degradation. Therefore, we added the associated reactions to the models corresponding to these 31 organisms. The remaining strains, namely *B. boum* DSM 20432*, B. coryneforme* DSM 20216*, B. thermacidophilum subsp. thermacidophilum* DSM 15837*, B. pseudolongum subsp. pseudolongum* DSM 20099 and *B. ruminantium* DSM 6489 lack the corresponding enzymes.

In summary, all the models were expanded, and validated by comparing with previously known substrate utilisation data for the exact strains. For strains that lack specific experimental observation on carbon source utilisation, we assumed the phenotype based on information from other strains of the same species included in this study. In addition, we performed BLAST to confirm for the presence of the enzymes associated with substrate utilisation. Furthermore, in the conditions where we lacked proper literature evidence on the substrates, we fell back upon the AGORA reconstruction but with our media constraints. A total of 21 different reactions were added across the 36 models, as shown in Supplementary Table 1. The exact modifications are detailed in Supplementary Table 2.

### Defining universal media for bifidobacterial growth

Initially, we generated a defined *in silico* medium for *B. bifidum* PRL 2010 based on the available experimental media components^42^. However, this defined set of *in silico* medium components failed to enable growth across all the 36 strains of *Bifidobacterium* considered here. The universal common set was generated by creating a union of all uptake components across strains. Each of these uptake components was removed from each strain and optimised for growth. Finally, we generated an augmented set of universal components, which can produce all biomass components across strains with glucose as the representative carbohydrate source (Supplementary Table 1).

### Model simulations

We simulated the models using the popular constraint-based analysis techniques, Flux Balance Analysis (FBA)^44,45^ and Flux Variability Analysis (FVA)^46^. FBA and FVA were used to analyse the individual capabilities of the strains, allowing uptake of a single nutrient source with all defined universal media components. We included 30 different carbon sources/nutrient environments encompassing mono-, di-, tri-, and polysaccharides (Supplementary Table 1). For the simulation of growth on different nutrient environments, the lower bound of the carbon uptake was set as 10 mmol/gDW/h, and a flux uptake of 1 mmol/gDW/h^26^ was used for each of the defined media components. For single species/strain metabolite production simulations, each of the models was first optimised for growth, and subsequently for metabolite production (acetate, lactate, ethanol, formate, and succinate). For metabolite production, the biomass reaction was constrained to be equal to the maximum growth rate achieved. For a given carbon source, we considered a threshold growth rate of > 0.01 h^-1^ for defining an organism’s growth.

In addition, we performed *in silico* single reaction knockouts by setting the bounds of the corresponding reaction(s) to zero. We identified single lethals and synthetic lethal pairs across each of the strains of bifidobacteria, using the Fast-SL algorithm previously developed in our laboratory^47^, in 30 different environments (corresponding to the different carbon sources considered in this study) as well as in a rich media environment. The rich media mimic the environment where *Bifidobacterium* colonises, namely the large intestine which has abundant sources of complex carbohydrates. In rich media, all exchange reactions corresponding to 30 different carbon sources were constrained to 10 mmol/gDW/h. All simulations were performed on MATLAB 2016a (Mathworks Inc., USA) using the COBRA Toolbox^36^ and *Gurobi6* (Gurobi Optimization LLC, USA) as the solver for solving the optimisation problems corresponding to FBA and FVA.

### Clustering the bifidobacterial strains

To classify the bifidobacterial strains in order to better understand their metabolic capabilities and their role in the gut microbiome, we clustered them based on their growth profiles, SCFA production, and essential reactions. For each organism, we simulated the presence/absence of growth in each of the environments, and represented these observations as a binary vector (0 representing no growth, and 1 representing growth). We hierarchically clustered the organisms, using the average linkage algorithm as implemented in MATLAB clustergram, based on the Hamming distance between these binary vectors. This distance essentially captures how similar the growth *profile* of a pair of organisms is. Using the same information, we also clustered the SCFAs and single lethals, again using Hamming distance as the distance metric. We generated “clustergram” and dendrogram plots illustrating the clustering obtained.

### Data Availability

All data generated and analysed in this study are available in the supplementary information files.

## Acknowledgements

This work was supported by the Institute Post-Doctoral Fellowship, Indian Institute of Technology Madras to N. T. Devika.

## Contributions

N.T.D. and K.R. designed the study. N.T.D. performed the analyses. N.T.D. and K.R. analysed the data. N.T.D. prepared the original draft of the manuscript. N.T.D. and K.R. revised, read and approved the final manuscript.

## SUPPLEMENTARY INFORMATION

**Supplementary Fig 1:**
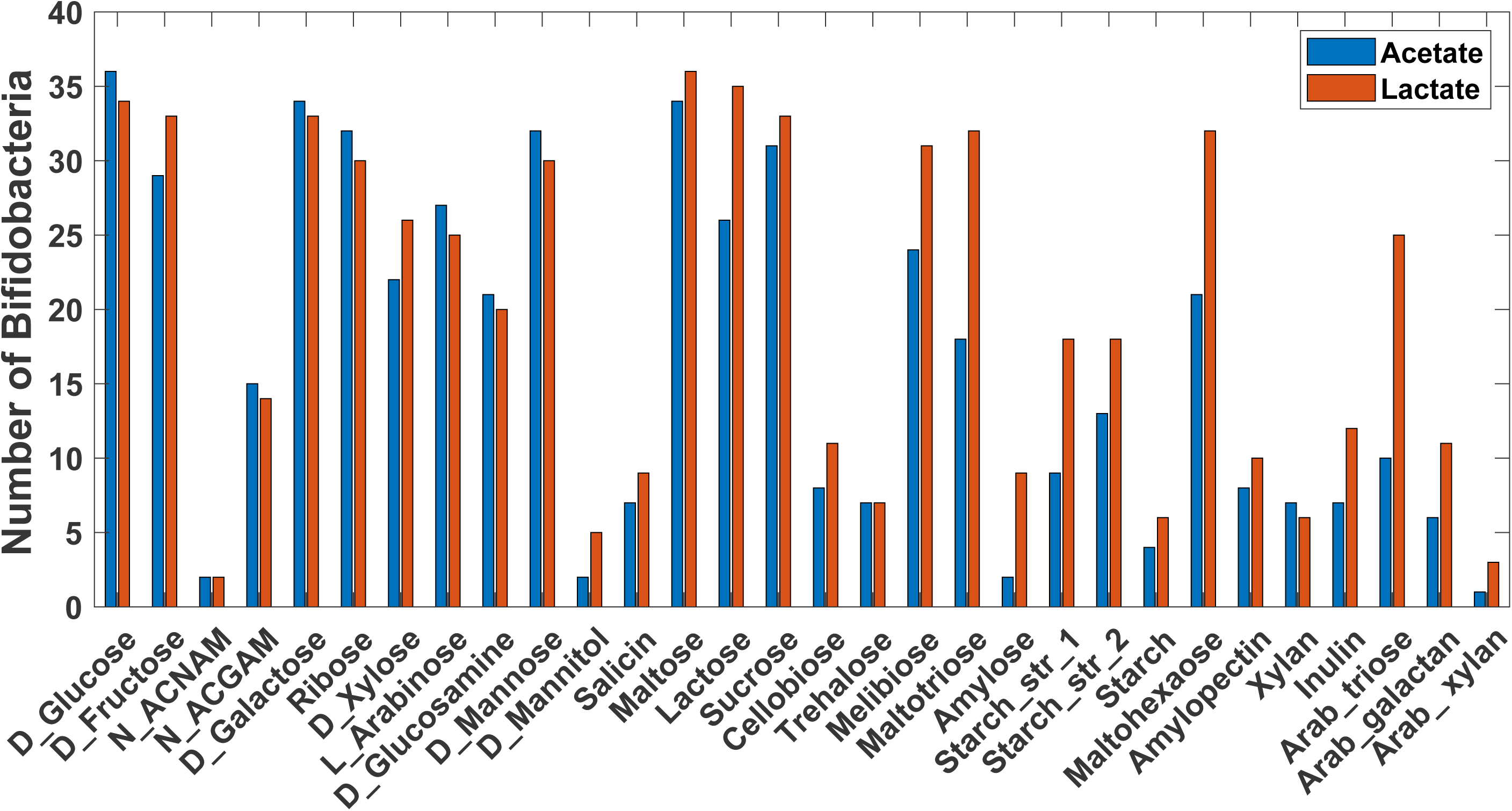
Environment dependent production of acetate and lactate across strains of bifidobacteria. Comparison of acetate and lactate production across strains of bifidobacteria under 30 different nutrient environments.

**Supplementary Table 1:** Description of organism, media components, carbon sources, reaction additions and knock-outs performed in the study.

**Supplementary Table 2:** Comparison of phenotype predictions between AGORA and our enhanced (re-curated) models with literature references.

**Supplementary Table 3:** List of single and double lethals computed across strains of bifidobacteria using Fast-SL in rich environment.

